# A *nadA* mutation confers nicotinic acid auxotrophy in pro-carcinogenic intestinal *Escherichia coli* NC101

**DOI:** 10.1101/2021.02.12.431052

**Authors:** Lacey R. Lopez, Cassandra J. Barlogio, Christopher A. Broberg, Jeremy Wang, Janelle C. Arthur

**Author notes:** Locus Biosciences, Inc., Durham, North Carolina, United States of America (Cassandra J. Barlogio). Department of Chemistry, The University of North Carolina at Chapel Hill, Chapel Hill, North Carolina, United States of America (Christopher A. Broberg). Address correspondence to Janelle C. Arthur.

## Abstract

Inflammatory bowel diseases and inflammation-associated colorectal cancer are linked to blooms of adherent-invasive *Escherichia coli* (AIEC) in the intestinal microbiota. AIEC are functionally defined by their ability to adhere/invade epithelial cells and survive/replicate within macrophages. Changes in micronutrient availability can alter AIEC physiology and interactions with host cells. Thus, culturing AIEC for mechanistic investigations often involves precise nutrient formulation. We observed that the pro-inflammatory and pro-carcinogenic AIEC strain NC101 failed to grow in minimal media (MM). We hypothesized that NC101 was unable to synthesize a vital micronutrient normally found in the host gut. Through nutrient supplementation studies, we identified that NC101 is a nicotinic acid (NA) auxotroph. NA auxotrophy was not observed in the other non-toxigenic *E. coli* or AIEC strains we tested. Sequencing revealed NC101 has a missense mutation in *nadA*, a gene encoding quinolinate synthase A that is important for *de novo* NAD biosynthesis. Correcting the identified *nadA* point mutation restored NC101 prototrophy without impacting AIEC function, including motility and AIEC-defining survival in macrophages. Our findings, along with the generation of a prototrophic NC101 strain, will greatly enhance the ability to perform *in vitro* functional studies that are needed for mechanistic investigations on the role of intestinal *E. coli* in digestive disease.

**Importance:** Inflammatory bowel diseases (IBD) and colorectal cancer (CRC) are significant global health concerns that are influenced by gut resident microbes, like adherent-invasive *Escherichia coli* (AIEC). Nutrient availability influences specialized metabolite production, AIEC-defining functional attributes, and AIEC:host interactions. NC101 is a pro-inflammatory and pro-carcinogenic AIEC strain commonly used for studies on IBD and CRC. We identified that NC101 growth *in vitro* requires a micronutrient found in the host gut. By correcting an identified mutation, we generated an NC101 strain that no longer has micronutrient restrictions. Our findings will facilitate future research that necessitates precise nutrient manipulation, enhancing AIEC functional studies and investigations on other auxotrophic intestinal microbiota members. Broadly, this will improve the study of bacterial:host interactions impacting health and disease.

## Introduction

Inflammatory bowel diseases (IBD), including Crohn’s disease and ulcerative colitis, are a major global health concern that affects over 3 million adults in the United States alone (1, 2). IBD is a chronic and multifactorial disease that is driven by aberrant immune responses to commensal microbes, genetic susceptibility, and environmental factors (3). IBD patients experience painful, chronic, and relapsing intestinal inflammation that can lead to life-threatening complications, including intestinal fibrosis and colorectal cancer (CRC) (4, 5). Experimental models have demonstrated that IBD and CRC can be driven by the intestinal microbiota and that specific microbes, such as *Escherichia coli*, are associated with human disease (6, 7). IBD and CRC have no single etiology and no cure (8, 9). Therefore, understanding the function of disease-associated gut microbes may uncover novel therapeutic options for intestinal diseases, like IBD and CRC.

Intestinal microbes influence the onset and progression of IBD and CRC via metabolite production and modulation of mucosal immunity (10–14). *E. coli* are common inhabitants of the intestinal microbiota (15, 16). Strain level differences can alter the pro-inflammatory or pro-carcinogenic potential of *E. coli*, partly through changes in small molecule production (12–14, 17, 18). A pathovar of *E. coli*, termed adherent-invasive *E. coli* (AIEC), are enriched in the gut microbiota of human IBD and CRC patients (19). AIEC exacerbate experimental colitis and promote CRC in a variety of murine models (17, 18, 20–24). There is no genetic definition for AIEC (14, 19, 25). Instead, AIEC are classically defined by their ability to adhere/invade epithelial cells and survive/replicate within macrophages (19, 26). Environmental conditions, including nutrient availability and intestinal inflammation, can alter AIEC behavior and impact intestinal colonization and disease (14, 17, 27–30). Therefore, the ability to precisely manipulate AIEC growth conditions is essential for *in vitro* studies investigating AIEC behavior and production of pro-inflammatory and pro-carcinogenic molecules.

*E. coli* NC101 is a well-known AIEC strain utilized by numerous investigators to study how intestinal *E. coli* adapt to and influence the host during IBD and CRC (14, 17, 18, 24, 31–33). NC101 was originally isolated from a specific pathogen free wild-type mouse at North Carolina State University (34). Colonizing wild-type mice with NC101 does not induce intestinal pathology, even during monoassociation studies using gnotobiotic animals (17). However, despite a lack of traditional toxins and virulence factors, NC101 induces antigen-driven intestinal inflammation in genetically-susceptible IBD mouse models (e.g. interleukin 10 deficient mice) (34). Thus, NC101 is considered a pathobiont and a highly relevant model organism for defining how susceptible individuals may mount inappropriate immune responses to seemingly innocuous intestinal *E. coli*.

NC101 adapts to the inflamed intestinal milieu by modulating expression of its gene repertoire (35, 36). Nutrient availability alters AIEC physiology, persistence in the microbiota, and production of pro-inflammatory and pro-carcinogenic mediators (14, 17, 27–30). Monoassociation studies with gnotobiotic mice have led to the discovery of several AIEC-derived host-influencing molecules (i.e. specialized metabolites) that drive inflammation and tumorigenesis, including yersiniabactin and colibactin (17–19). Like many specialized metabolites, yersiniabactin and colibactin are produced via biosynthetic gene clusters that can be activated by changes in micronutrient availability, notably iron (37, 38). The nature of AIEC-derived specialized metabolites makes them difficult to isolate and study in functional assays. Therefore, the repertoire of AIEC-derived metabolites and their impact on the host has been largely unexplored.

Variations in micronutrient availability can impact the virulence and physiology of AIEC (27, 28, 39). Therefore, culturing AIEC for mechanistic studies necessitates using a simplified base media that allows for precise nutrient manipulation. During our studies, we observed that modified M9 minimal media (MM) does not sustain NC101 growth *in vitro*. We hypothesized that NC101 was an auxotroph. Through nutrient supplementation studies, we discovered that NC101 requires nicotinic acid (NA, niacin, Vitamin B3) for growth. NA auxotrophy was not observed in other non-toxigenic laboratory *E. coli* strains (K12 or 25922), AIEC, or non-AIEC human intestinal strains (40). Genetic evaluation revealed that NC101 has a missense mutation in the NAD biosynthesis gene (*nadA*) that encodes for quinolinate synthase A. Importantly, we generated a prototrophic NC101 revertant strain that eliminated *E. coli* micronutrient restraints. Correcting NC101 auxotrophy had negligible impact on NC101 function, including motility and AIEC-defining survival in macrophages.

NC101 micronutrient constraints have limited our ability to perform *in vitro* functional studies, which often require careful nutrient manipulation. Overall, our findings will enable precise nutrient manipulation for mechanistic studies on auxotrophic microbiota members, like AIEC, *Shigella spp*., or Uropathogenic *E. coli* (41–44). Importantly, our work will facilitate *in vitro* functional assays and small molecule purification efforts with the pro-inflammatory and pro-carcinogenic AIEC strain NC101. Furthermore, these studies will broadly improve our understanding of the microbiota in intestinal diseases like IBD and CRC.

## Results

### The pro-carcinogenic adherent-invasive *E. coli* strain NC101 requires nicotinic acid to sustain growth

During *in vitro* studies to evaluate AIEC function in long-term culture (24+ hr), we attempted to passage NC101 in modified M9 minimal media (MM) that includes glycerol and casamino acids. NC101 can successfully be subcultured from Luria-Bertani (LB) agar or broth, a rich medium, to MM (17, 28). However, NC101 failed to grow when subcultured from MM to MM (**Fig. 1A-C**). We hypothesized that NC101 was an auxotroph, unable to synthesize a key nutrient found in the murine gut. *Shigella spp*., a transient gut pathogen and close relative of *E. coli*, are generally nicotinic acid (NA, Niacin, Vitamin B3) auxotrophs (41, 43–45). Thus, we specifically tested whether vitamin supplementation could restore NC101 growth in MM. Supplementing MM with a complex Vitamin Mix (VM) restored NC101 growth at 8hr and 24hr (**Fig. 1A-B**).

**Fig. 1.**
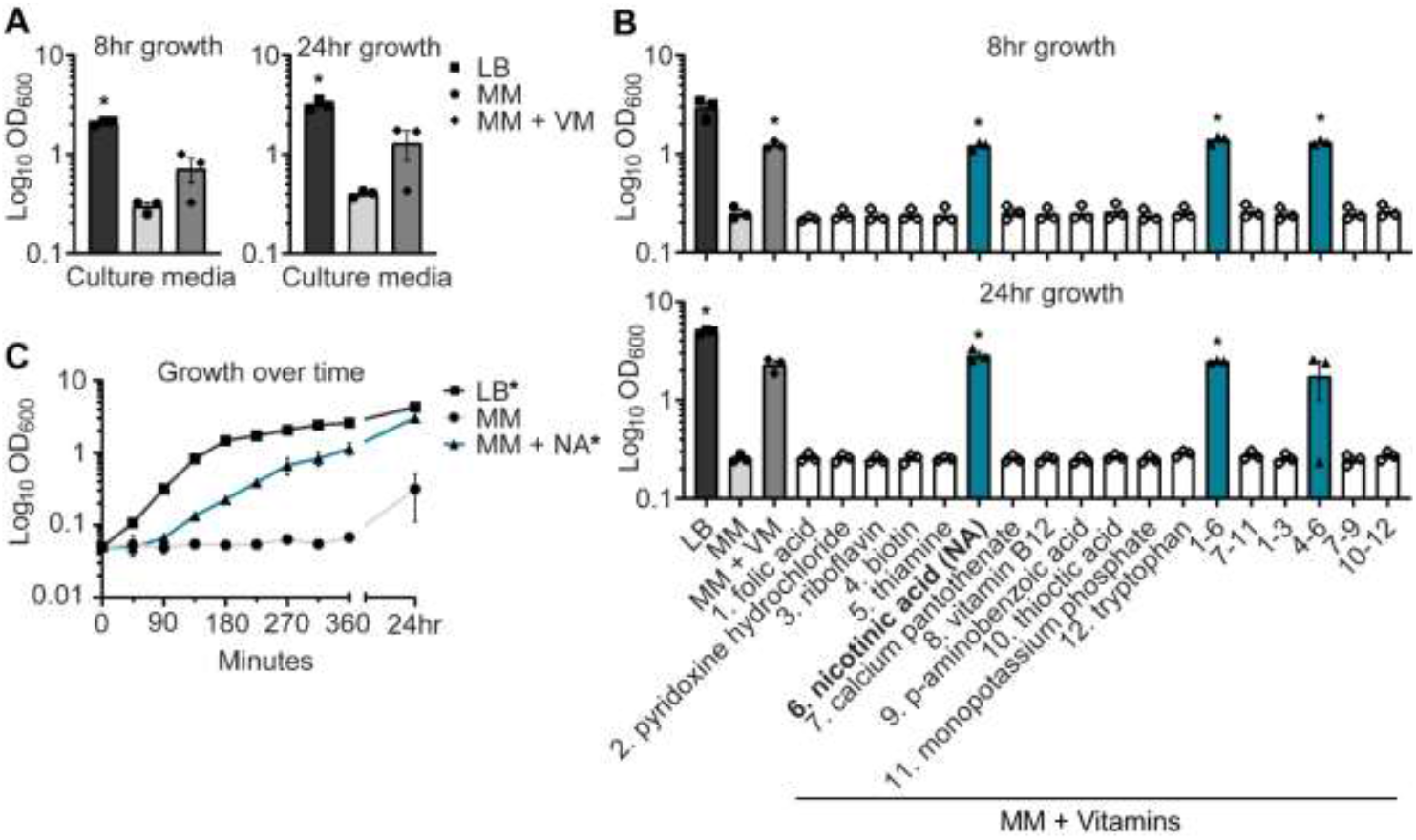
Nicotinic acid restores growth of *E. coli* NC101 in minimal media. (**A**) Wild-type NC101 was grown in Luria-Bertani (LB) broth, minimal media (MM), or MM + vitamin mix (VM). Growth was measured at 8hr and 24hr by culture optical density, OD_600_. (**B**) NC101 was grown in LB, MM, MM + VM, or MM supplemented with individual or combinations of vitamins and tryptophan. OD_600_ was assessed at 8hr and 24hr. (**C**) Growth curve of NC101 in LB, MM, or MM + NA. (**A, B**) Bars (n = 3) or (**C**) points (n = 4) depict mean +/- SEM. Significance (*) is shown compared to NC101 growth in MM at (**A,B**) each timepoint or (**C**) 24hr and was determined at p < 0.05, using a one-way ANOVA with Dunnett’s T3 multiple comparisons test.

To identify which vitamin(s) in the VM were essential for NC101 growth, we supplemented MM with individual or combinations of VM components and assessed NC101 growth. Tryptophan supplementation was also tested, as tryptophan metabolism can be influenced by host-microbe interactions in the gut (46). Only MM containing NA, alone or in combination, sustained NC101 growth in MM at 8hr and 24hr (**Fig. 1B**). Further, NA alone restored normal NC101 growth kinetics in MM and significantly enhanced growth at 24hr (**Fig. 1C**). NC101 grew when subcultured from MM to LB, indicating NC101 does not have a global growth defect (**Fig. 1A-C**). Together, these data suggest NC101 is an NA auxotroph.

### NA auxotrophy is not a defining feature of non-toxigenic *E. coli*

Resident non-toxigenic *E. coli* are common among the intestinal microbiota and many are considered commensal strains (15, 16). Yet, other *E. coli* (e.g. AIEC) are associated with chronic intestinal inflammation and may be referred to as pathobionts (15, 19). We questioned whether NA auxotrophy was shared across clinically derived non-toxigenic *E. coli*. In addition to evaluating model *E. coli* strains (K12 and 25922), we evaluated clinical specimens isolated from the intestinal mucosa of IBD or non-IBD patients (*E. coli* LF82, 42ET-1, 568-3, HM670, 37RT-2, 532-9, and 39ES-1) (40, 47, 48) (**Table 1**). These clinical isolates have been characterized in the lab from which they originated for AIEC status, and at least partial genome sequences are available for all strains (40). To determine the extent of NA-dependency among these strains, we passaged isolates in MM with and without NA and assessed growth by measuring optical density (OD_600_) at 2hr, 4hr, 8hr, and 24hr (**Fig. 2A**). We again observed that NC101 had a growth defect in MM, detectable at 2hr and continuing through 24hr (**Fig. 2A**). However, all examined laboratory strains and clinical isolates grew in MM with and without NA. (**Fig 2A**). Therefore, NA auxotrophy does not appear to be a defining characteristic shared by non-toxigenic resident intestinal *E. coli*.

**Table 1.** *Escherichia coli* strains used in this study.

| Strain | Description of non-virulent<br>*E. coli* strains | Isolated<br>from: | Adherent-invasive *E. coli*<br>(AIEC) Status | Reference<br>or Source |
| --- | --- | --- | --- | --- |
| Wild-type<br>(WT) NC101 | Streptomycin resistant (Str<sup>R</sup>)<br>isolate of classical murine-<br>derived AIEC | Laboratory<br>*E. coli* isolate | AIEC | (34), This<br>study |
| LF82 | Classical human-adapted<br>clinical AIEC isolate | Crohn's<br>disease<br>patient | AIEC | (48) |
| 42ET-1 | Clinical AIEC isolate | Non-IBD<sup>a</sup><br>patient | AIEC | (40) |
| 568-3 | Clinical AIEC isolate | Crohn's<br>disease<br>patient | AIEC | (40) |
| K12<br>(MG1655) | Model *E. coli* strain | Laboratory<br>*E. coli* isolate | Non-AIEC | ATCC<sup>®</sup><br>700926<sup>TM</sup> |
| 25922 | Model *E. coli* strain | Patient | Non-AIEC | ATCC<sup>®</sup><br>25922<sup>TM</sup> |
| HM670 | Clinical *E. coli* isolate | Crohn's<br>disease<br>patient | Non-AIEC, but<br>has enhanced<br>survival in<br>macrophages | (47) |
| 37RT-2 | Clinical *E. coli* isolate | Non-IBD<sup>a</sup><br>patient | Non-AIEC | (40) |
| 532-9 | Clinical *E. coli* isolate | Crohn's<br>disease<br>patient | Non-AIEC | (40) |
| 39ES-1 | Clinical *E. coli* isolate | Crohn's<br>disease<br>patient | Non-AIEC | (40) |
| Nicotinic<br>acid revertant<br>(NA<sub>derivative</sub><br>or NA<sub>D</sub>)<br>NC101 | Spontaneous prototrophic<br>revertant of WT NC101<br>(G263T mutation in *nadA*) | Laboratory<br>*E. coli* isolate | N.D.<sup>b</sup>, but<br>exhibits survival<br>in macrophages | This study |
| NC101 Δ*fliC* | Non-motile flagellin mutant<br>derived from WT NC101 | Laboratory<br>*E. coli* isolate | N.D.<sup>b</sup> | This study |

<sup>a</sup>IBD = inflammatory bowel disease, <sup>b</sup>N.D. = not determined

**Fig. 2.**
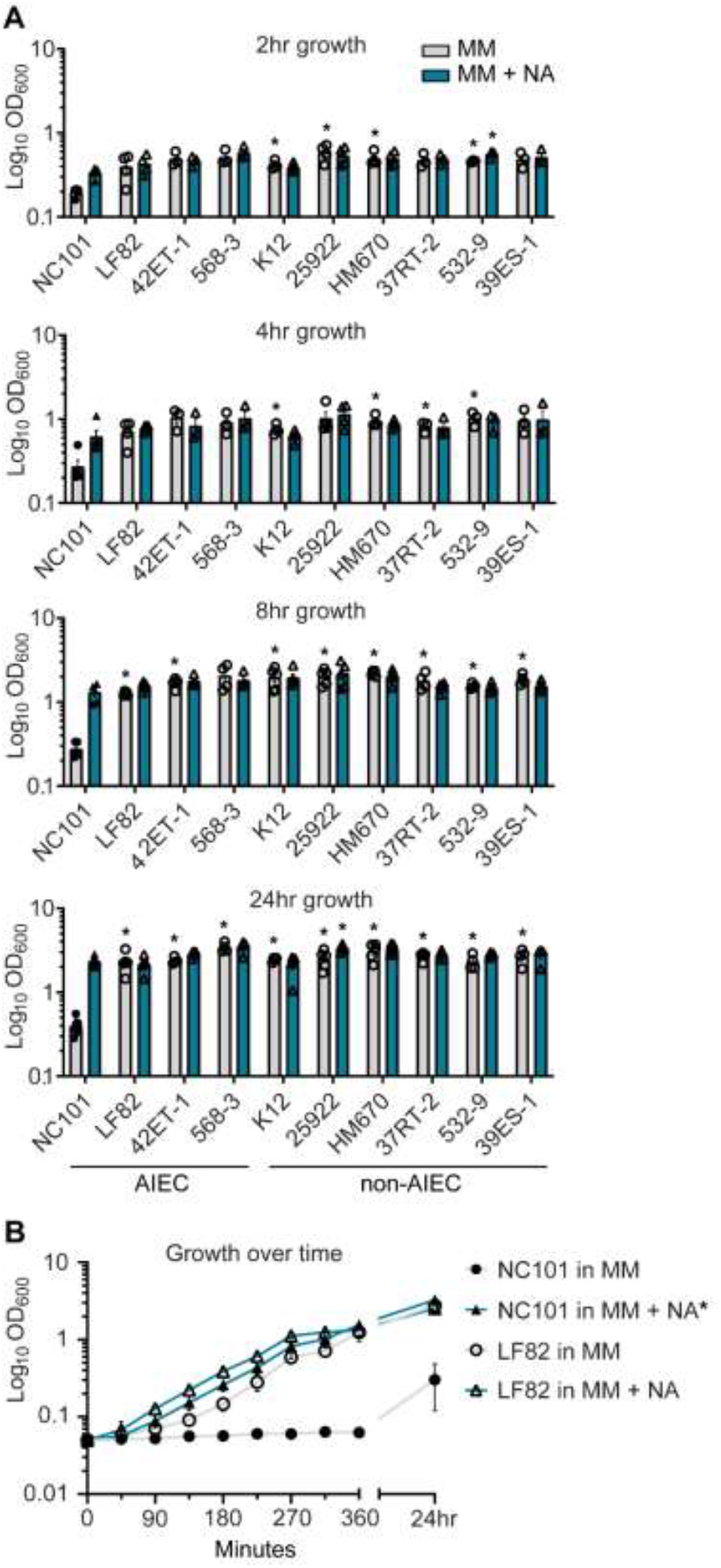
NA auxotrophy is not common among non-toxigenic *E. coli* strains, including prototypic AIEC LF82. (**A**) AIEC and non-AIEC *E. coli* isolates, were grown in minimal media (MM) with and without nicotinic acid (NA). Growth was evaluated at 2hr, 4hr, 8hr, and 24hr by OD_600_. (**B**) Growth curve of wild-type NC101 and LF82 (human-derived AIEC) in MM with and without NA. (**A**) Bars (n = 3-6) or (**B**) points (n = 3) depict mean +/- SEM. (**A**) Strains grown in MM were compared to NC101 grown in MM, and strains grown in MM + NA were compared to NC101 grown in MM + NA. Significance (*) is shown compared at (**A**) each timepoint or (**B**) 24hr and was determined at p < 0.05, using a one-way ANOVA with Dunnett’s T3 multiple comparisons test.

LF82 is a well-known human-derived AIEC strain that can grow in MM without NA (48) (**Fig. 2A-B**). We directly compared the growth kinetics of NC101 and LF82 in MM with and without NA (**Fig. 2B**). While early growth of LF82 in MM was minimally enhanced by NA, this difference was indistinguishable by 6hr (**Fig. 2B**). Thus, the prototypic AIEC strain LF82 does not exhibit NA auxotrophy. Combined with our findings in Fig. 2A, we conclude that NA auxotrophy is not an AIEC-defining feature.

### NC101 has a defect in the *de novo* NAD biosynthesis pathway

NA is a precursor for NAD biosynthesis (41). NAD is an electron carrier and an essential cofactor for bacterial metabolism (41). In *E. coli* and related bacteria, NAD can be synthesized *de novo* from L-aspartate (L-asp) through the generation of quinolinic acid (quinolinate, Qa). In this process, Quinolinate synthase A and B (encoded by *nadA* and *nadB*, respectively) catalyze the oxidation of L-asp to iminoaspartate and condensation with dihydroxyacetone phosphate to generate quinolinate (41). Quinolinate is converted to nicotinic mononucleotide (NaMN) by a *nadC* encoded enzyme and ultimately NAD via enzymes encoded by *nadD* and *nadE* (41). NAD biosynthesis can also occur through salvage pathways that utilize vitamin precursors like NA or nicotinamide (Nm) (41, 43, 49) (**Fig. 3A**).

**Fig. 3.**
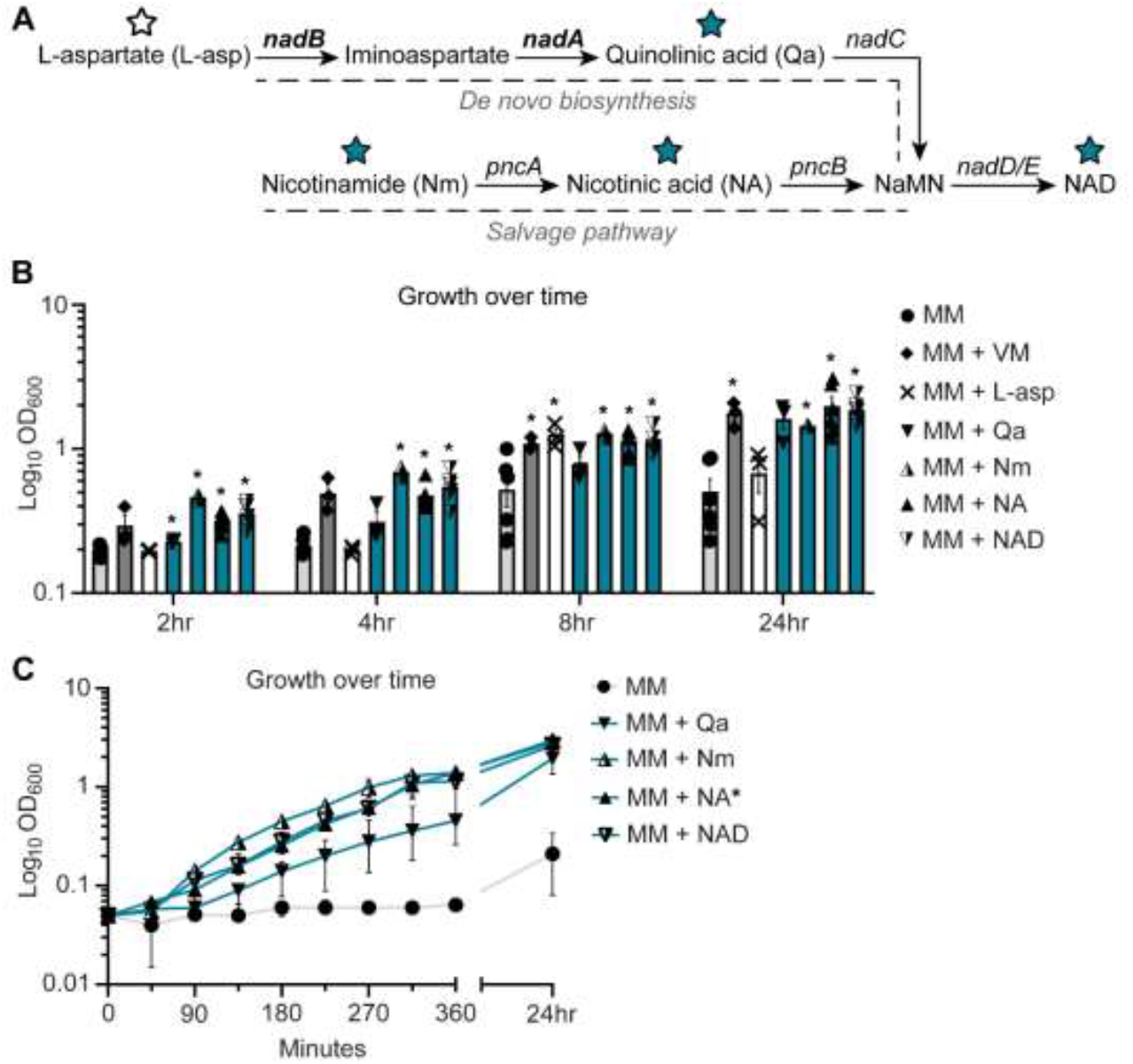
NC101 has a defect in the *de novo* NAD biosynthesis pathway. (**A**) Illustration of NAD biosynthesis pathway in *E. coli*, including pathway intermediates and genes involved. Intermediates tested in *E. coli* growth assays are indicated by stars (blue restored growth, white did not). Bolded genes, *nadA* and *nadB*, are predicted to be responsible for NC101 auxotrophy. (**B**) Wild-type NC101 was grown in minimal media (MM), MM + vitamin mix (VM), or MM + NAD biosynthesis pathway intermediates: L-aspartate (L-Asp), quinolinic acid (Qa), nicotinamide (Nm), nicotinic acid (NA), and NAD. Culture density was evaluated at 2hr, 4hr, 8hr, or 24hr by OD_600_. (**C**) Growth curve of NC101 in MM with or without Qa, Nm, NA, or NAD. (**B**) Bars (n = 3-6) or (**C**) points (n = 3) depict mean +/- SEM. Significance (*) is shown compared to NC101 growth in MM at (**B**) each timepoint or (**C**) 24hr and was determined at p < 0.05, using a one-way ANOVA with Dunnett’s T3 multiple comparisons test.

Since NA restored the growth of NC101, we predicted that NC101 had a defect within the NAD biosynthesis pathway. To determine whether this was the case, we assessed NC101 growth in MM supplemented with key NAD biosynthesis intermediates: L-asp, Qa, Nm, NA, and NAD. L-asp failed to consistently sustain NC101 growth in MM. Conversely, Qa sustained NC101 growth in MM and Nm, NA, and NAD significantly restored growth across all timepoints (**Fig. 3B**). Growth curves revealed the kinetics of enhanced NC101 growth in the presence of the restorative NAD biosynthesis intermediates: Qa, Nm, Na, and NAD (**Fig. 3C**). When examining the NAD biosynthesis pathway, this indicated that NC101 likely had a defect in the NAD biosynthesis genes *nadA* or *nadB* (**Fig. 3A**)

### NA auxotrophy in NC101 is linked to a mutation in NAD biosynthesis gene *nadA*

After our growth supplementation assays revealed a likely defect in *nadA* or *nadB*, we sought to identify the genetic factor(s) responsible for NA auxotrophy in NC101. We performed whole genome sequencing on wild-type (WT) NC101 and compared the sequence to prototrophic *E. coli*, LF82 and K12. Sequencing revealed that WT NC101 has a missense mutation in *nadA* (T263G) that was associated with auxotrophy (**Fig. 4A**).

**Fig. 4.**
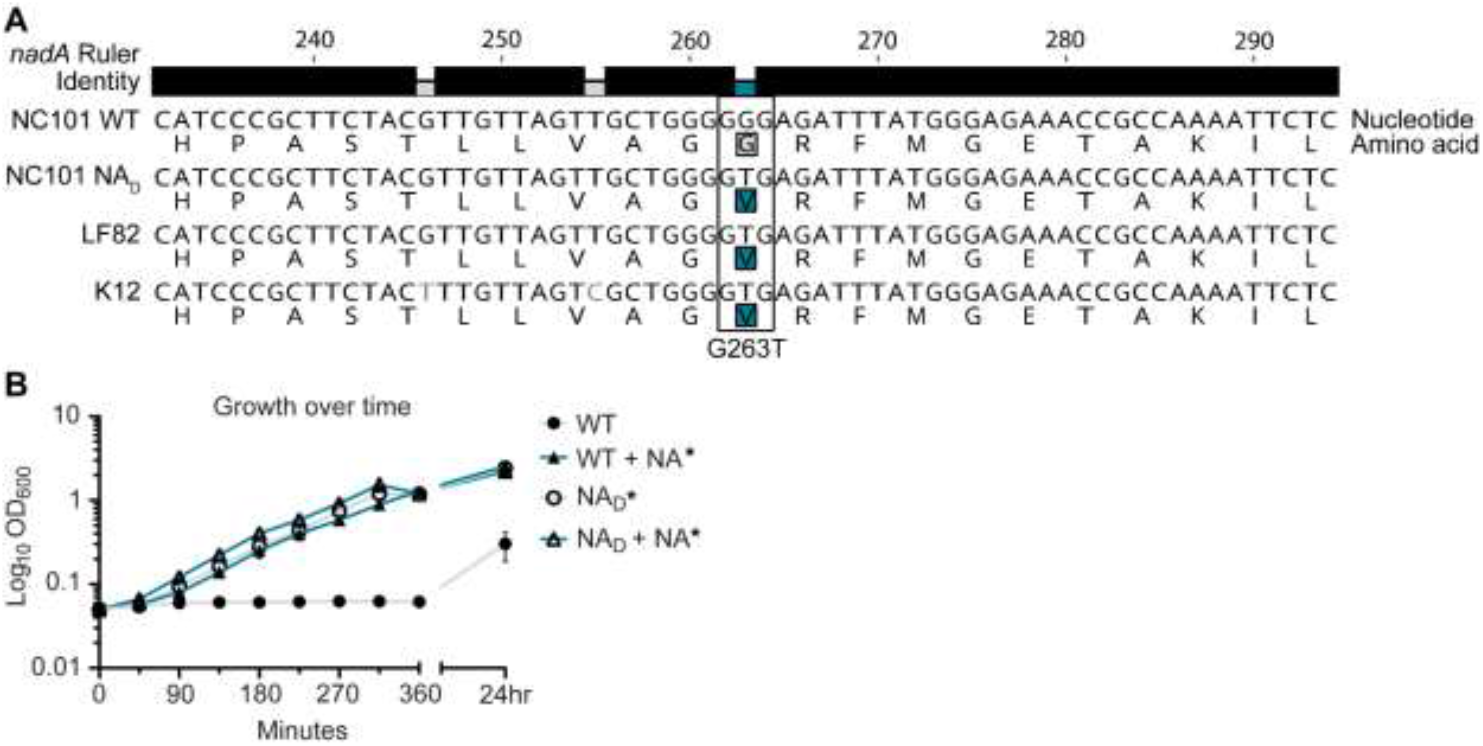
A missense mutation in NAD biosynthesis gene *nadA* confers NA auxotrophy in NC101. (**A**) Genetic alignment of partial *nadA* sequence from NA auxotrophic (Wild-type (WT) NC101) and prototrophic (NC101 NA_D_, LF82, and K12) *E. coli*. Nucleotide and amino acid sequences, noted by 1 letter abbreviations, are shown. The ruler displays nucleotide position of coding sequence. The identity bar displays regions of similarity (black) or dissimilarity (grey or blue). The highlighted amino acids show the region of noted dissimilarity (*nadA* G263T) between NA auxotrophic (grey) and prototrophic (blue) *E. coli*. (**B**) A growth curve of WT NC101 and prototrophic revertant NC101 strain (NA_D_) in minimal media (MM) with and without nicotinic acid (NA). Growth was measured by culture optical density, OD_600_. Points depict mean +/- SEM (n = 4). Significance (*) is shown compared to NC101 growth in MM at 24hr and was determined at p < 0.05, using a one-way ANOVA with Dunnett’s T3 multiple comparisons test.

To further validate the genetic determinants of NC101 NA auxotrophy, we generated a prototrophic strain by passaging WT NC101 on MM agar plates in the absence of NA (42). Sequencing of a selected spontaneous prototrophic revertant, termed NA_Derivative_ or NA_D_ NC101, revealed that NA_D_ NC101 had a single nucleotide substitution in *nadA* (G263T, compared to WT) that matched the prototrophic *E. coli* strains LF82 and K12 (**Fig. 4A**). It is important to note that NA_D_ NC101 also had a silent mutation in an intergenic region that was absent from WT NC101, but we predict this mutation had no impact on NA_D_ NC101 prototrophy (Accession #SAMN16810912) (**Table 2**). To support that NA auxotrophy is due to the observed *nadA* mutation, our whole genome sequencing revealed that two other NC101 spontaneous prototrophic revertants had missense mutations in *nadA* – one of which shared the same *nadA* (G263T, compared to WT) nucleotide substitution as NA_D_ NC101 (Accession #SAMN16810913 and #SAMN16810914) (**Table 2**).

**Table 2.**
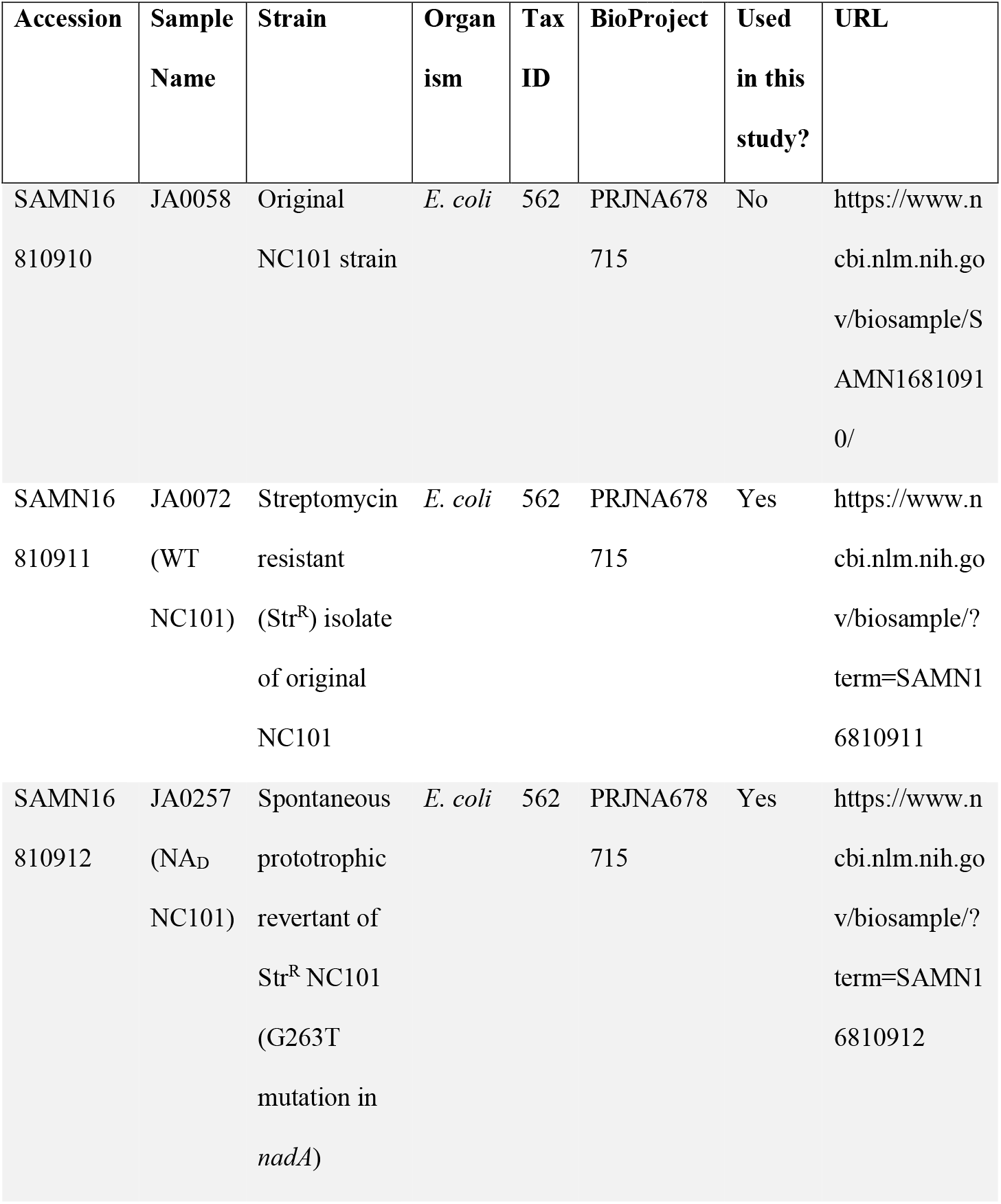
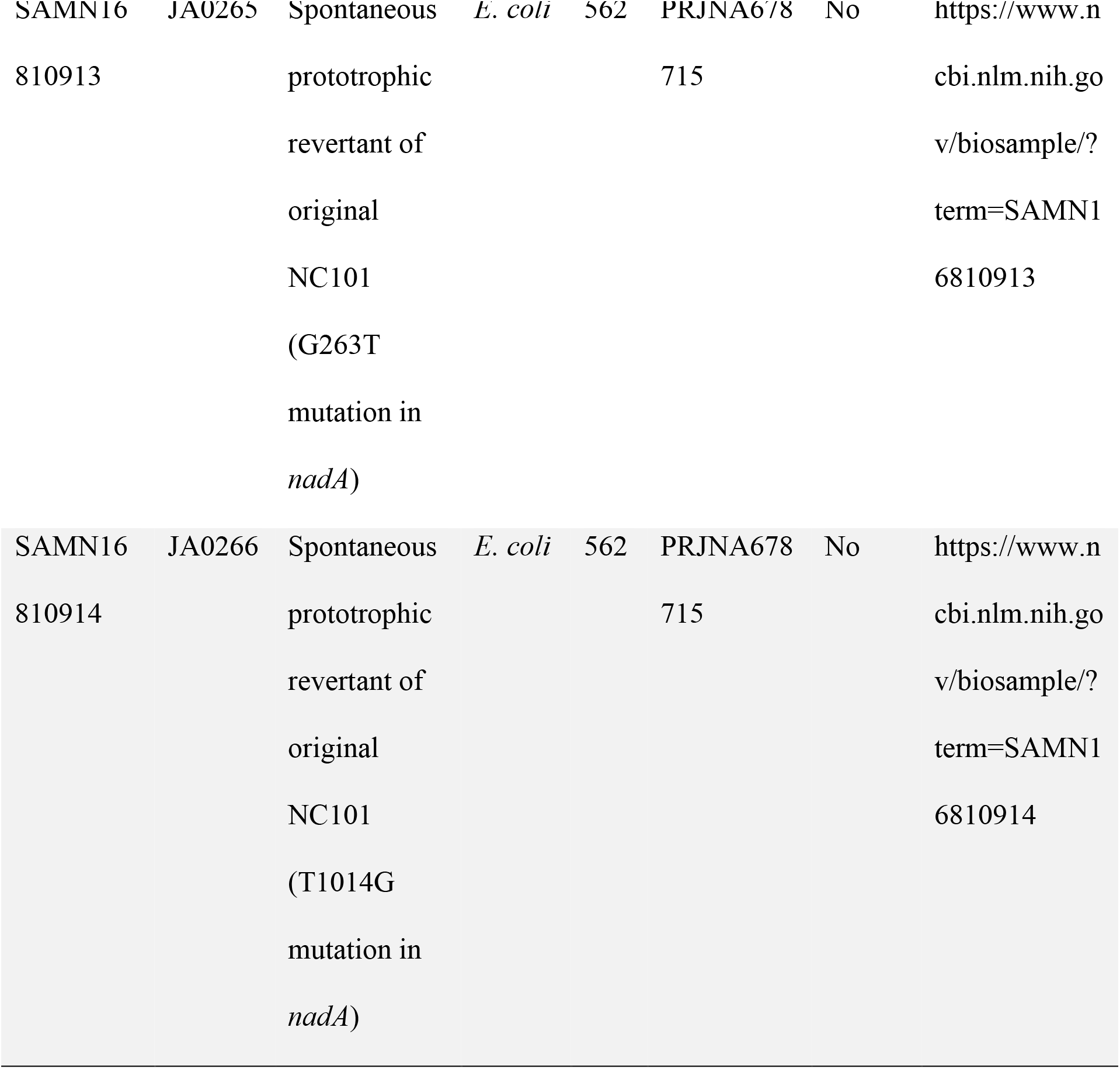
Repository information for published genomic sequences.

To confirm that the NA_D_ revertant restored prototrophy, we grew WT NC101 and NA_D_ in MM with and without NA. NA_D_ grew significantly better in MM versus WT NC101 at 24hr (**Fig. 4B**). Growth of NA_D_ in the absence of NA was the equivalent to WT grown with added NA, as noted by overlapping growth curves (**Fig. 4B**). The addition of NA did not significantly enhance NA_D_ growth in MM, suggesting NA auxotrophy was successfully eliminated in this strain (**Fig. 4B**). These findings are consistent with the literature, which indicates NadA is important for NAD biosynthesis and mutations in *nadA* can drive NA auxotrophy in *E. coli, Shigella spp*., and *Salmonella spp*. (41, 43, 44, 50). Therefore, our data support that NC101 NA auxotrophy is due to a mutation in *nadA*.

### Correcting NA auxotrophy in NC101 has negligible impact on bacterial motility or AIEC-associated survival in macrophages

To determine if correcting NA auxotrophy impacted AIEC physiology and interactions with mammalian cells, we assessed WT and NA_D_ NC101 for motility and survival in macrophages. Motility is not an AIEC-defining feature, but hypermotility has recently been linked to changes in AIEC:host interactions (19, 51). Due to the WT NA auxotrophy, motility was only assessed on MM agar plates supplemented with NA. There was no significant difference in motility between WT and NA_D_ NC101 in the presence of NA (**Fig. 5A**). However, the motility of both WT and NA_D_ NC101 differed significantly from the non-motile control mutant, NC101 Δ*fliC* (flagellar filament structural protein) (51) (**Fig. 5A**).

**Fig. 5.**
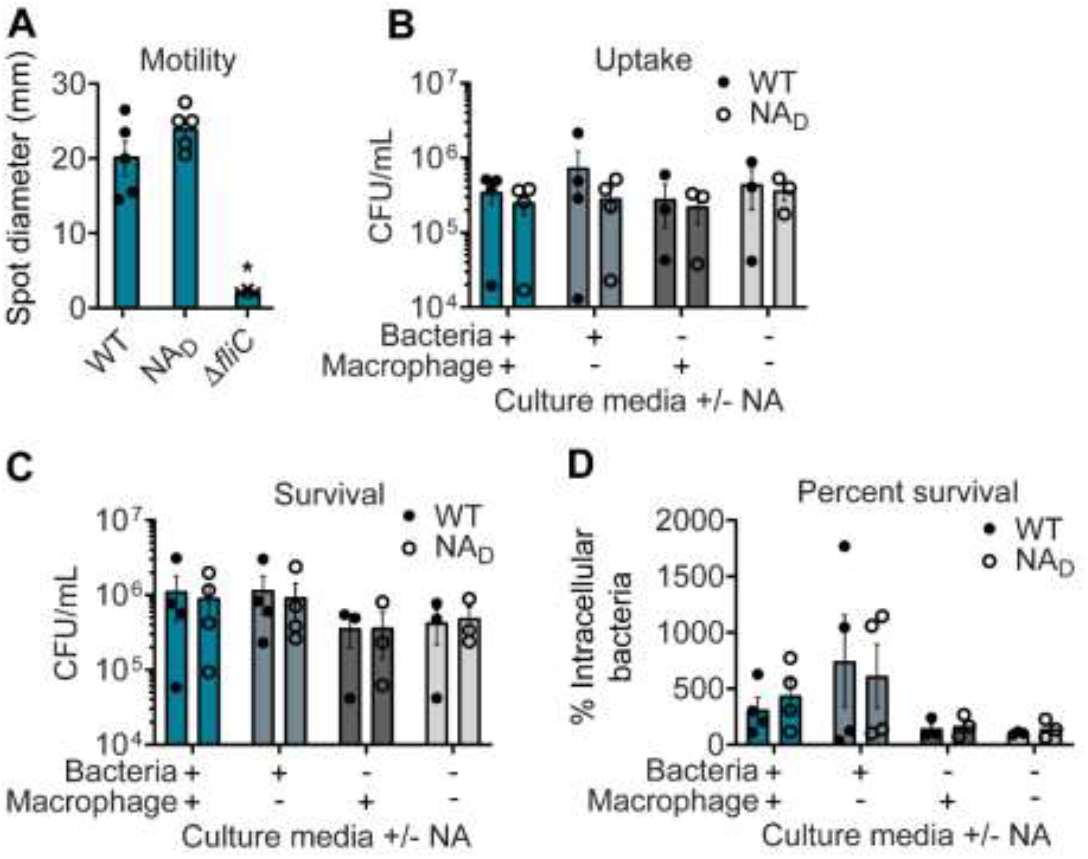
Correcting NA auxotrophy in NC101 has minimal impact on *in vitro* AIEC function. (**A**) Wild-type (WT) NC101, prototrophic NC101 NA_D_, and non-motile control NC101 Δ*fliC* were grown on minimal media (MM) soft agar plates with nicotinic acid (NA). Diameter of motility swarm spots (mm) were measured at 8hr (n = 3-5). (**B-D**) J774A.1 murine macrophages were infected at a multiplicity of infection (MOI) = 10 with WT or NA_D_. Bacterial culture media before infection (“Bacteria”) or cell culture media during infection (“Macrophage”) were with or without NA supplementation. Number of bacteria are shown at (**B**) 1hr or (**C**) 24hr post-infection as Log10 colony forming units (CFU)/mL. (**D**) Percent survival = [(CFU/mL at 24hr)/(CFU/mL at 1hr]×100 (n = 3-4). Bars depict mean +/- SEM. Significance (*) is shown compared to WT NC101 and was determined at p < 0.05, using a (**A**) one-way ANOVA with Dunnett’s T3 multiple comparisons test or (**B-D**) Welch’s t test.

A key feature of AIEC is enhanced survival in macrophages, a characteristic linked to their pro-inflammatory activities (26, 52). To evaluate whether the identified *nadA* mutation impacted AIEC-defining survival in macrophages, WT and NA_D_ NC101 were subcultured in MM with and without NA and used to infect macrophage cell cultures, which were also maintained in the presence or absence of NA. Despite the expected differences in culture densities between WT and NA_D_ strains upon subculturing in MM without NA (**Fig. 1C**), we could obtain a sufficient amount of WT NC101 to infect with an equivalent multiplicity of infection for all experiments. Baseline macrophage cell culture media contains an excess of NA, so as expected, there were not WT NC101 survival defects in the infection assay. Importantly, we found there were no significant differences in AIEC intramacrophage uptake (1hr) or survival (24hr) between WT and NA_D_ NC101 in the presence or absence of NA supplementation (**Fig 5B-D**). Therefore, eliminating NA auxotrophy in NA_D_ NC101 had negligible impact on these AIEC-associated functions.

## Discussion

*E. coli* are common members of the mammalian microbiota (15, 16). Many *E. coli* isolates are prototrophic (41). However, a study identified that NA auxotrophy was common among the B2 phylotype of *E. coli* strains that are usual intestinal inhabitants (41). Herein, we illustrate that the AIEC strain NC101 (phylotype B2) is an NA auxotroph due to a missense mutation in NAD biosynthesis gene *nadA*. These findings are significant, as NC101 is an established AIEC often used for studies on IBD and CRC; yet, NA auxotrophy in NC101 has not been defined (14, 17, 18, 24, 31–33).

It is unclear why NC101 possesses NA auxotrophy versus the human-derived resident *E. coli* we examined (**Fig. 2**). Perhaps some feature of the murine intestinal environment promoted this NC101 characteristic. Genome reduction or loss-of-function mutations may facilitate adaptation to the intestinal microenvironment, as the decrease in biosynthetic cost of compounds likely provides an advantage when key nutrients are consistently present within an environment (53, 54). It is possible that loss of NAD biosynthesis gene function represents a way in which AIEC NC101 adapted to survive within the murine host. For example, an abundance of NA in the murine diet may have permitted murine-adapted NC101 with a mutation in *nadA* to persist in the gut, despite NA auxotrophy. It is also possible that among the many stochastic mutations experienced by *E. coli* strains, this *nadA* mutation simply conferred no benefit or detriment, allowing it to persist as a resident microbe of the murine gastrointestinal tract.

Besides reducing biosynthetic cost, Na/NAD play a key role in virulence and signaling across various microbial species, including *E. coli, Shigella spp.,Candida glabrata, Bordetella pertussis, and Legionella pneumophila* (41–44, 55–58) In *E. coli*, NA can regulate the EvgA/EvgS two-component regulatory system that drives multidrug resistance and acid tolerance (56, 59). While in *Shigella spp*., a pathogen but close relative of nonpathogenic *E. coli*, loss of functional NAD biosynthesis genes (often *nadA* and/or *nadB*) reduces *Shigella* virulence and alters interactions with host cells (44, 55). However, our results demonstrate that NA auxotrophy does not impact a pro-inflammatory and key defining feature of AIEC, survival in macrophages.

The intestinal microbiota comprises a large/diverse collection of host-associated microbes, microbial genes, and products (6). Our lab and others have been interested in pro-inflammatory and pro-carcinogenic molecules derived from intestinal bacteria, namely yersiniabactin and colibactin (17, 18). Many host-influencing microbial-derived molecules, often termed specialized metabolites, are produced by sophisticated multi-enzymatic machinery encoded by bacterial biosynthetic gene clusters. By nature, many of these specialized metabolites are difficult to isolate and purify in sufficient amounts for functional analysis. However, their production can often be activated by nutrient deficiency (12, 39, 60). Therefore, studies on specialized metabolites and their interactions with host cells often requires precise nutrient manipulation to study *in vitro*. To optimize the production of these unique bioactive molecules and reduce non-essential media components that complicate purification, we have identified the minimal media components necessary to grow the model AIEC NC101 and generated an NC101 strain no longer restricted by NA auxotrophy. This strain, NC101 NA_D_, can easily be cultured in MM for functional studies or used to purify AIEC specialized metabolites. Ultimately this strain, NA_D_ NC101, can now serve as a research tool to investigate how precise nutrient manipulation impacts AIEC behavior under minimal media conditions.

In summary, our work in defining and correcting the NA auxotrophy in AIEC NC101 will 1) enable precise nutrient manipulation for *in vitro* studies on AIEC as they relate to IBD and CRC, 2) inform culture-based methods to evaluate the function of other auxotrophic gut microbiota members and their metabolites, and 3) facilitate small molecule isolation and purification from the pro-inflammatory and pro-carcinogenic strain NC101. Long-term, we expect our findings will contribute to the identification of microbiota-derived prognostic and therapeutic targets for human digestive diseases.

## Materials and Methods

### Bacterial strains

Descriptions of *E. coli* strains used in this study are listed in **Table 1**. NC101 Δ*fliC* was generated using the λ-red recombinase method, as previously described (17, 61). Primers used for Δ*fliC* generation are listed in **Table 3.**

**Table 3.**
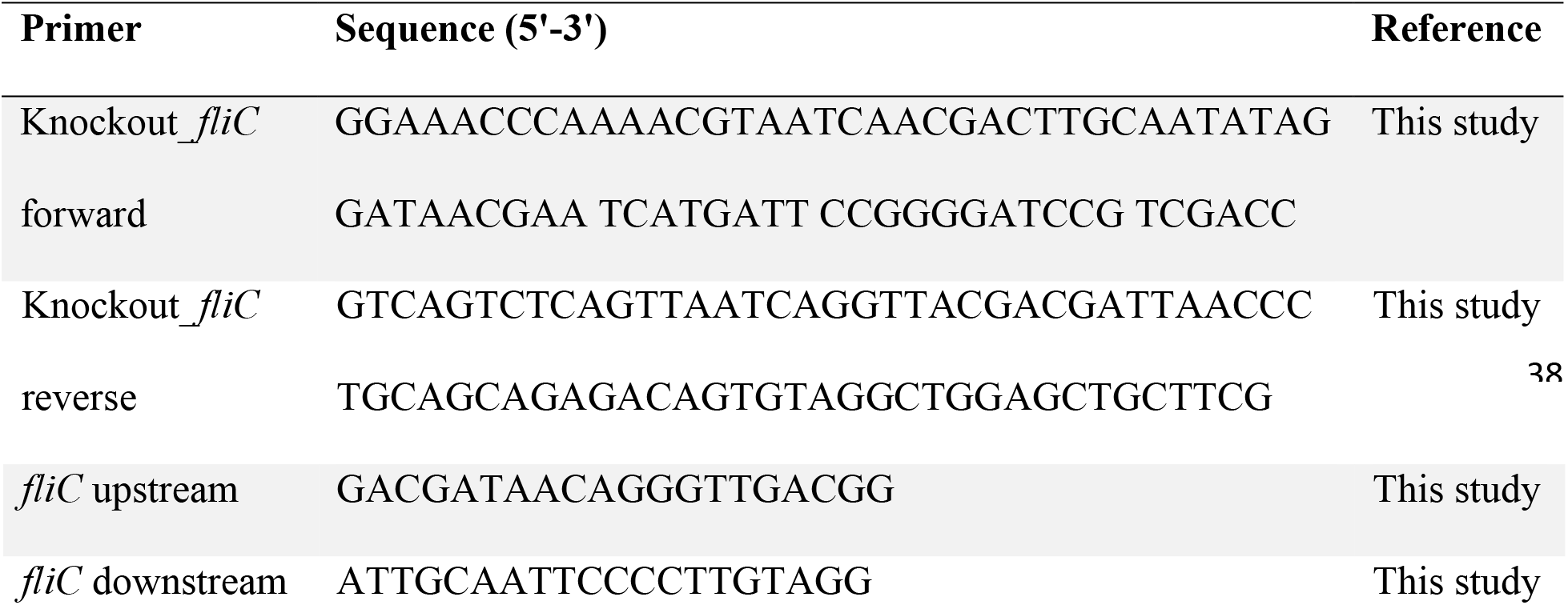
Primers used for strain construction.

### Media composition

#### M9 minimal-defined media (MM) – 5X M9 salts

64g Na_2_HPO_4_*7H_2_O, 15g anhydrous KH_2_PO4, 2.5g NaCl, and 5g NH_4_Cl brought to 1L in diH_2_O (62). **Complete MM:** 0.1mM CaCl_2_, 1X M9 salts, 2mM MgSO_4_, 0.4% glycerol, and 0.2% casamino acids (CAA, Sigma #2240) brought to 1L in diH_2_O. Where indicated, the following were added at these final concentrations: nicotinic acid (NA, Sigma #N4126), 50μg/L; L-aspartate (L-asp), 200mg/L; nicotinamide (Nm), 50μg/L; and NAD, 1μg/L.

#### Vitamin mix (VM), 100X stock

2mg folic acid, 10mg pyridoxine hydrochloride, 5mg riboflavin, 2mg biotin, 5mg thiamine, 5mg nicotinic acid, 5mg calcium pantothenate, 0.1mg vitamin B12, 5mg p-aminobenzoic acid, 5mg thioctic acid, and 900mg monopotassium phosphate brought to 1L in diH_2_O and aliquoted into 10mL stocks. One 10mL stock was used per liter of media. Formulation is from ATCC and is based on Wolfe’s Vitamin solution (ATCC^®^ MD-VS^™^).

#### Overnight cultures

Bacterial strains were preserved at ^-^80°C and grown overnight at 37°C on Luria Bertani (LB, Fisher Sc. #BP9722-2) agar plates. Isolated colonies were transferred to MM and grown overnight (>15hrs) at 37°C with shaking at 220 rpm.

#### Growth assays

Overnight cultures were centrifuged and washed three times with 1X phosphate buffered saline (PBS), to remove any trace compounds contained in the culture. Cells were re-suspended and normalized by optical density (OD_600_) in test media. Cultures were grown at 37°C with shaking at 220 rpm. For passaging assays, OD_600_ was recorded at the indicated timepoints (2hr, 4hr, 8hr or 24hr). For growth curves, OD_600_ was recorded every 45min for 6hr and a final timepoint was recorded at 24hr.

#### Spontaneous prototrophic revertant generation

WT NC101 was grown overnight in MM + NA, 5mL of the culture was centrifuged, and the supernatant discarded. The cell pellet was washed twice with 1X PBS and resuspended in 500μL 1X PBS. A 100 μl spot was spread onto each of five MM agar plates without NA. Plates were incubated at 37°C and monitored for growth of revertant colonies (42). Colonies were grown on MM without NA to confirm prototrophy and isolates were preserved at −80°C. Whole genome sequencing was performed to determine the location and nature of the mutation(s) leading to reversion. The revertant used in these studies was termed NC101 NADerivative (NA_D_).

#### NC101 genome assembly

A complete NC101 genome was assembled from nanopore sequence using Minimap2 and Miniasm (63). The assembly was circularized and polished four times with Racon (64) followed by once with Medaka (Oxford Nanopore Technologies, https://github.com/nanoporetech/medaka). Matched Illumina sequence data was used to polish the resulting assembly using FMLRC (65) with parameters “-k 21 -K 30 -m 3 -f 0.05 -B 10”. The final polished genome was rotated and linearized such that it starts at the origin of replication.

#### Whole genome sequencing

Three spontaneous prototrophic revertants, including NA_D_, were sent for whole genome sequencing. Samples were sent to the Microbial Genome Sequencing Center (MiGS), formerly at the University of Pittsburgh, for genomic DNA extraction and Illumina 2×150 paired end sequencing on the NextSeq 550 platform. Sequencing reads were mapped to our closed NC101 genome using CLC Genomic Workbench7.5.1 with average coverage of 85x for JA0257, 62x for JA0265, and 75x for JA0266. Sequences for LF82 (NC_011993.1) and K12 (NC_000913.3) were obtained from the National Center for Biotechnology Information (NCBI) and all alignments were analyzed via Geneious Prime version 2020.1.2. Assembled sequences from this study were deposited in NCBI and repository information is listed in **Table 2**.

#### Motility

Isolates were grown overnight, as described above. A 1μL spot was used to inoculate the center of MM soft agar plates (MM + 0.25% agar) with NA. Plates were incubated at 37°C for 8hr and the diameters of motility swarms were measured.

#### Macrophage survival assays

Bacterial intramacrophage survival was measured using the standard gentamicin protection assay for AIEC bacteria (26, 28). The J774A.1 murine macrophage-like cell line was used as a model and maintained according to ATCC standards in DMEM + 10% heat-inactivated FBS (DMEM, Gibco #11995-065). J774A.1 cells were seeded at 2×10^5^ cells/mL in 1mL media into 24-well plates (Falcon #353047) and grown overnight. The next day, bacterial overnight cultures were subcultured in MM with and without NA for 3hr. Before infection, J774A. 1 monolayers were washed twice with 1X PBS. Then, subcultured bacteria were added at a multiplicity of infection (MOI) = 10 in cell culture media with and without NA. Plates were spun at 180 × g for 5min. Prepared bacterial cultures were serial diluted and plated on LB agar plates to validate infection dose.

After a 30min incubation at 37°C with 5% CO_2_, infected cultures were washed twice with 1X PBS and gentamicin-laden media was added (100μg/mL gentamicin for 1hr timepoint and 20μg/mL for 24hr in DMEM + 10% FBS with and without NA). At 1hr and 24hr, cells were washed twice with 1X PBS and 500μL of 1% Triton X-100 in diH_2_O was added to each well for 5min. Samples were mixed, serial diluted, and plated on LB agar plates to determine viable colony forming units (CFU). Percent intracellular bacteria = [(CFU/mL at 24hr)/(CFU/mL at 1hr]×100.

#### Statistics

Statistical analysis was performed using Prism version 9.0.0 (GraphPad software San Diego, CA). A Welch’s t test was used when two experimental groups were compared and a one-way ANOVA with Dunnett’s T3 multiple comparisons test was used when 3 or more experimental groups were compared. Differences with a p-value less than 0.05 were considered significant. All experiments included at least 3 biological replicates with 1-2 technical replicates each, per timepoint.

#### Data availability

Assembled sequences from our whole genome sequencing, above, were deposited in NCBI and repository information is listed in **Table 2**.

## Acknowledgments

We thank the following colleagues for kindly gifting *E. coli* strains: R. Balfour Sartor, University of North Carolina (UNC) at Chapel Hill (*E. coli* NC101, LF82, and K12 MG1655); Kenneth W. Simpson, Cornell University (*E. coli* 42ET-1, 568-3, 37RT-2, 532-9, and 39ES-1); Leslie M. Hicks, UNC at Chapel Hill (*E. coli* 25922); Barry J. Campbell, University of Liverpool (*E. coli* HM670); Melissa Ellermann, University of South Carolina at Columbia (NC101 *AfliC)*. We thank Dr. Kimberly Walker for critical reading of the manuscript. NC101 genome assembly was supported by Jeremy Wang NIH K01DK119582 (JW). The remaining body of this work was supported by funding from NIH R01DK124617 (JCA), pilot funding from NIH P30DK034987 (subcontract JCA), and a UNC Lineberger Comprehensive Cancer Center Innovation Award (JCA).

